# Small-molecule-based regulation of gene expression in human astrocytes upon both positive and negative actuations of G-quadruplex control systems

**DOI:** 10.1101/2024.02.16.580621

**Authors:** M. J. Vijay Kumar, Jérémie Mitteaux, Zi Wang, Ellery Wheeler, Nitin Tandon, Sung Yun Jung, Robert H. E. Hudson, David Monchaud, Andrey S. Tsvetkov

## Abstract

A great deal of attention is currently being paid to strategies aiming at uncovering the biology of the four-stranded nucleic acid structure G-quadruplex (G4) *vi*a their stabilization in cells using G4-specific ligands. The conventional definition of chemical biology implies that a complete assessment of G4 biology can only be achieved through the implementation of a complementary approach involving the destabilization of cellular G4s by *ad hoc* molecular effectors. We report here on an unprecedented comparison of the cellular consequences of G4 chemical stabilization by pyridostatin (PDS) and destabilization by phenylpyrrolocytosine (PhpC) at both transcriptome- and proteome-wide scales in patient-derived human brain cells. Our results show that the stabilization of G4s by PDS triggers the dysregulation of many cellular circuitries, the most drastic effects originating in the downregulation of 354 transcripts and 158 proteins primarily involved in RNA transactions. In contrast, the destabilization of G4s by PhpC modulates the G4 landscapes in a far more focused manner with the upregulation of 295 proteins mostly involved in RNA transactions as well, thus mirroring the effects of PDS. Our study is the first of its kind to report on the extent of G4-associated cellular circuitries in human cells by systematically pitting the effect of G4 stabilization against destabilization in a direct and unbiased manner.

## Introduction

Chemical biology^1^ hinges on the strategic use of small molecule modulators to disrupt cellular equilibria reversibly, both spatially and temporally.^2^ The responses of cells to these molecular effectors are then analyzed to gain insights into underlying molecular mechanisms. For accurate and dependable interpretations, interventions in chemical biology rely on exquisitely specific molecular tools to prevent off-target effects and potential misinterpretations of molecular mechanisms. Ideally, chemical biologists should use both positive and negative regulators to ensure a balanced assessment of the cellular circuitry being studied, thereby enhancing the reliability of their findings.^3^

Recently, the field of DNA and RNA G-quadruplexes (G4s)^4–7^ has become a favorite playground for chemical biologists^8^ eager to uncover a new level of regulations of gene expression referred to as G4omics.^9, 10^ G4 is a four-stranded nucleic acid structure (**Figure 1A**) that assembles from guanine (G)-rich DNA or RNA sequences when freed from their hybridized forms (*e.g*., duplex for DNA, stem-loop for RNA).^11^ The G4-forming sequences are widespread in our genome and transcriptome, with >1 million putative G4 sequences predicted by G4Hunter (DNA)^12^ and G4RNA Screener (RNA).^13^ G4s fold thanks to the formation and self-stacking of cyclic arrays of Gs known as G-quartets; despite very high thermodynamic stability, they undergo rapid folding and unfolding orchestrated by G4 chaperones^14^ and G4 helicases,^15^ respectively, which intervene in virtually all DNA- and RNA-dependent processes. This explains why mutations in G4 helicase-coding genes are linked to multiple life-threatening diseases, including inheritable genetic diseases (*e.g*., WRN is mutated in Werner syndrome, RTEL1 in dyskeratosis congenita, XPD in xeroderma pigmentosum and Cockayne syndrome, etc.), cancers and aging phenotypes.^15–19^

**Figure 1.**
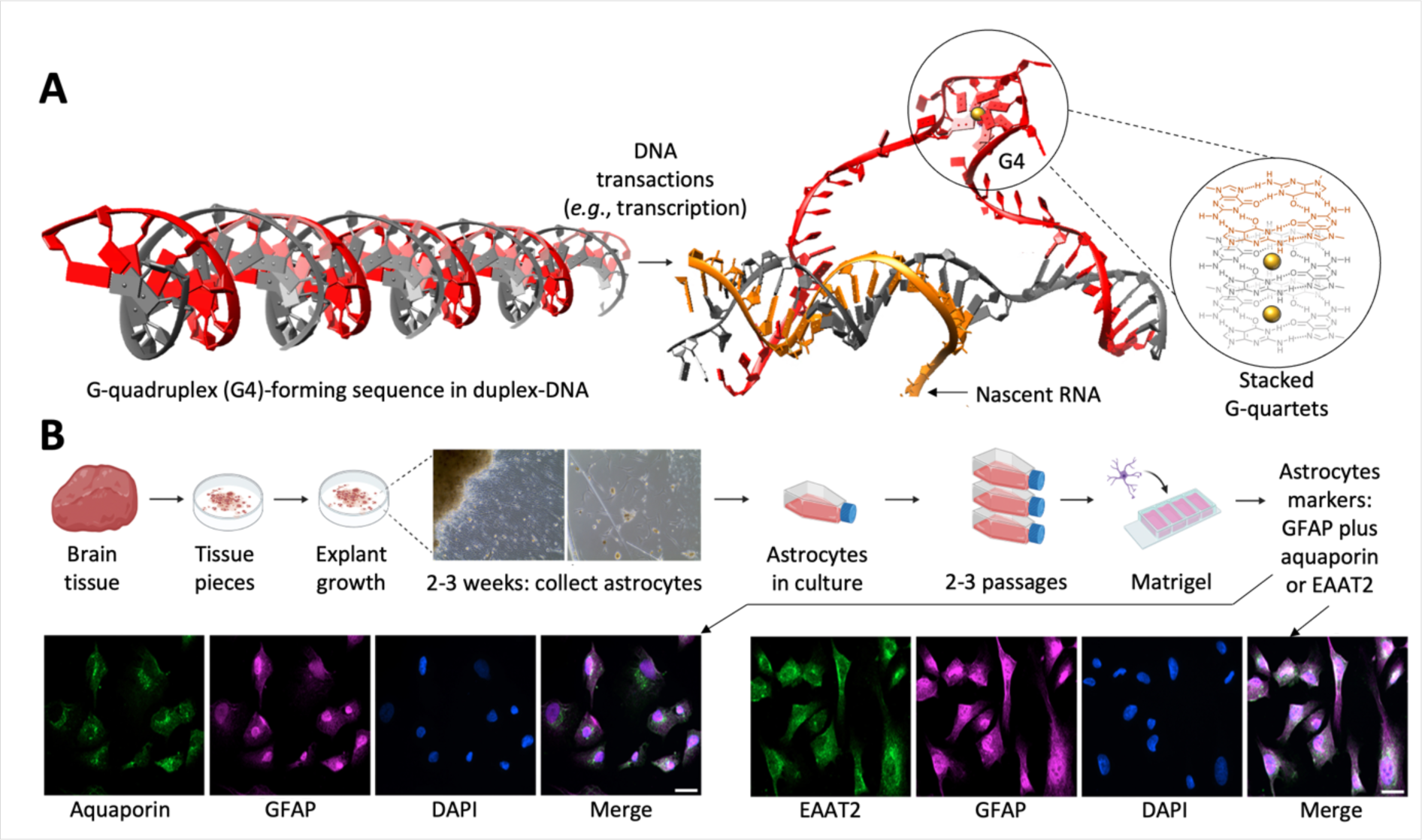
**A**. Schematic representation of the formation of a G4 structure originating in release of the G4-forming sequence from its duplex constraint (right panel: highlight on the self-stacking of G-quartets that forms the core of the G4 structure). **B**. Schematic representation of the collection and culture of astrocytes from human brain tissue (upper panel) and validation of their astrocytic phenotype by immunofluorescence (lower panel).

Chemical biologists aspire to substitute proteins with small molecules, which offer greater practical convenience. They have developed numerous G4 molecular chaperones, or G4 ligands,^20^ which have been used to *i-* demonstrate the existence and prevalence of both DNA and RNA G4s in human cells;^21^ *ii*-provide insights into the regulation of gene expression at both transcriptional and translational levels,^6, 22^ and *iii*-advance chemotherapeutic strategies that leverage G4s as genetic tools to impede the progression of diseases, especially cancers.^23^ In sharp contrast, the development of G4 molecular helicases, or G4 destabilizers, is still in its infancy:^17, 24^ While many G4 stabilizers have been used in yeast, cancer cells, and more recently in brains cells (*vide infra*), only a handful of G4 destabilizers have been developed so far,^25–30^ and their studies were limited to *in vitro* investigations. Only a phenylpyrrolocytosine referred to as PhpC^29^ has been identified as a reliable negative regulator of G4s in human cancer cells by both optical imaging and G4-RNA-specific affinity precipitation (G4RP) technique.^31, 32^

We aim here to assess the properties of PhpC in primary human astrocytes, major brain cells critically involved in age-associated brain disorders. Indeed, we recently provided evidence of the involvement of G4s in aging phenotypes, showing that *i*-aged mouse brains contain higher levels of G4s than young brains,^33^ *ii*-the stabilization of G4s with the small molecule stabilizer pyridostatin (PDS)^34^ promotes genomic instability in neurons, astrocytes, and microglial cells,^35–37^ and *iii*-mice injected with PDS exhibit cognitive impairment and accelerated aging,^33^ reminiscently of what was described in *C. elegans*.^38^ Aging indeed provides a permissive chromatin environment for G4s: beyond multiple epigenetic and epitranscriptomic changes, condensed heterochromatic regions are diminished during aging,^39–42^ with a frequent loss of nucleosomes, which provides a window of opportunity for G4-forming sequences to fold. G4s might thus play an important role in aging cells, but, at present, the associated molecular mechanisms are largely unknown.

The goal of the present study is to establish a firm connection between G4 stabilization and destabilization and nervous cell biology.^43–45^ To this end, our approach abides by the aforementioned definition of chemical biology, using both positive (PDS) and negative (PhpC) modulators of G4s in human astrocytes. We selected this cell type because astrocytes *i-* play a significant role in neuronal function, physiology, and overall brain development,^46^ *ii-* are well-accepted cell candidates for uncovering the pathophysiology of neurological disorders (considering their myriad roles in brain homeostasis), and *iii-* are practically convenient due to their ability to replicate and divide.^47–50^ We performed genome-wide gene expression analysis (RNA-seq) and proteome-wide mass spectrometry analysis to identify the molecular pathways affected by either PhpC or PDS. Our results show that G4s are indeed key genetic levers in astrocytes as the positive modulator PDS displays a strong inhibitory activity at both transcriptome- and proteome-wide scale, while the negative modulator PhpC affects the proteome of astrocytes only, positively contributing to the regulation of RNA processes, stress response, and cellular internal organization. These results thus show that G4s are interesting chemical biology targets in human astrocytes and that their destabilization by *ad hoc* molecular effectors might represent a relevant strategy to alleviate G4-associated dysfunctions.

## Results

Astrocytes were cultured from explants derived from the frontal lobes of human brain resections obtained from a 32-year-old patient. The astrocytic phenotype of these isolated cells was rigorously confirmed by immunostaining performed with a panel of well-established astrocyte-specific markers, including aquaporin,^51^ excitatory amino acid transporter 2 (EAAT2),^52^ glial fibrillary acidic protein (GFAP)^53^ S100β and vimentin^54^ (**Figure 1B**).

We first performed RNA-seq analysis for human astrocytes treated with 2 µM PDS for 24 h (**Figure 2**). The RNA-seq data demonstrated high data quality across all samples. The principal component analysis of this set of results highlighted clear distinctions between the untreated controls *versus* PDS-treated cells: we identified 516 (2.7%) differentially expressed genes (DEGs) out of a pool of 18,648 genes with measured expression. Among these 516 DEGs, 162 genes (31%) were upregulated, while 354 genes (69%) were downregulated (**Figure 2A** and Supplementary File 1). PDS treatment thus strongly alters the transcriptome of astrocytes, modulating the expression of an extended network of genes, in line with previous reports in different cell types.^37, 55^ We then conducted data mining within the context of both pathways (Kyoto Encyclopedia of Genes and Genomes (KEGG) database)^56^ and gene ontologies (Gene Ontology (GO) consortium database)^57, 58^ implementing iPathwayGuide (Advaita Bioinformatics),^59^ using the probability (*p*) value as a significance marker, considered to be high confidence when *p* ≤ 0.01. Our analysis revealed significant modifications encompassing 1,241 GO terms, 368 gene upstream regulators related to 28 pathways. The most enriched pathways comprised *i-* the regulation of cell cycle (31 differently expressed RNA out of 124 (31/124, 25%, including the cellular stress/DNA damage response protein GADD45G, *p* = 2.0e-3), *ii-* homologous recombination (9/41, 22%, DNA repair family (see the supplementary data for definition), including the helicase BLM and the recombinase RAD51, *p* = 1.0e-6), *iii-* DNA replication (12/36, 33%, including the helicase DNA2 (*p* = 4.3e-5) and the structure-specific endonuclease FEN1, *p* = 1.0e-6) and *iv-* the Fanconi anemia pathway (12/54, 22%, disease family, including the helicase FANCJ and the repair proteins BRCA1 and BRCA2, *p* = 1.0e-6) (**Figure 2C**). Among the most notable downregulated genes were thus helicases (20/72, 27%, including the G4 helicases BLM, FANCJ, and PIF1, *p* = 1.0e-6) and repair proteins (both double-strand break repair (39/246, 16%) and recombinational repair (30/139, 22%), including RAD51, BRCA1 and BRCA2, *p* = 1.0e-6, **Figure 2C**). These results make PDS a double-edged sword as it concomitantly inhibits G4 helicases, which thus triggers an increase in G4s, that is, in G4-mediated DNA damage,^60, 61^ and DNA repair proteins,^62^ which hampers the proper repair of this damage.

**Figure 2.**
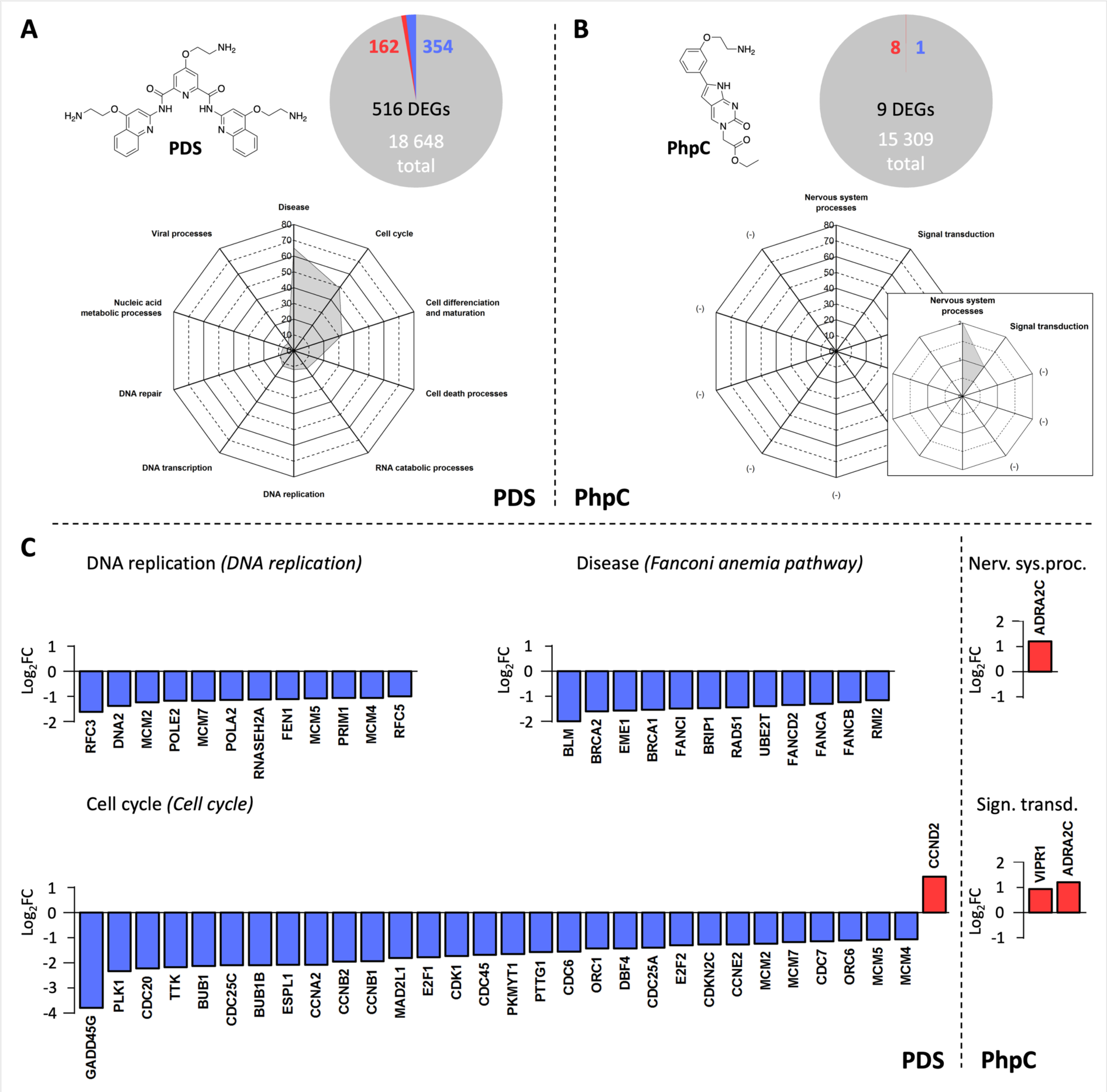
**A**. Chemical structure of pyridostatin (PDS), differentially expressed genes (DEGs) in human astrocytes upon incubation with PDS (pie chart) and dysregulated biological pathways (radar plot). **B**. Chemical structure of PhpC, DEGs upon incubation with PhpC (pie chart) and dysregulated biological pathways (radar plot). **C**. Examples of transcripts downregulated (blue) or upregulated (red) upon incubation with either PDS (left panel) or PhpC (right panel) belonging to families of dysregulated cellular pathways (see the supplementary data for definition).

In sharp contrast, PhpC-treated human astrocytes (20 μM for 24 h) experienced an almost insignificant transcriptomic change. We identified a total of only 9 DEGs (0.1%) out of a total of 15,309 genes with measured expression. Among these 9 DEGs, only 1 gene (11%) was downregulated, and 8 genes (89%) upregulated (**Figure 2B** and Supplementary File 1). The treatment of PhpC thus triggers the modification of only 286 GO terms, 5 gene upstream regulators (with moderate significance, *p* < 0.04), related to 5 pathways but in a minimal manner since the expression of only 1 to 4% of RNA (from 1/22 (hormone biosynthesis) to 2/129 RNA (ligand-receptor interaction)) were modified (again with moderate significance, *p* < 0.05, **Figure 2C**). Collectively, these results indicate that PhpC does not affect the transcription, likely because PhpC does not act on G4 DNA in the human astrocytes. We nevertheless compared the gene expression common between PDS- and PhpC-treated astrocytes and identified only 2 common DEGs (ADRA2C and ANKHD1-EIF4EBP3, *p* < 0.01), but not related to any recognized cellular pathway.

To investigate further the properties of these 2 molecular effectors, we focused on proteomics. Similar to the RNA-seq analysis, PDS-treated astrocytes experienced a series of significant changes compared to the untreated control (**Figure 3**). We identified 381 (10.6%) differentially expressed proteins (DEPs) out of a pool of 3,604 proteins with measured expression. Among these 381 DEPs, 223 proteins (58%) were upregulated, while 158 proteins (42%) were downregulated (**Figure 3A** and Supplementary File 2). Our study shows that PDS influences 417 GO terms, 57 upstream regulators related to 12 pathways. The most significantly impacted pathways include chromatin assembly (31 differently expressed proteins out of 80 (31/80, 39%)), nucleosome assembly (29/73, 40%), DNA packaging (31/92, 34%), all belonging to the chromatin modification and organization family, DNA conformation change (36/127, 28%), telomere organization (23/80, 29%), and ribonucleoprotein complex biogenesis (38/237, 16%) among others. These modifications include the dysregulation of the remodeler ATRX (*p* = 0.01, helicase family) as well as both the structure-specific endonuclease FEN1 (*p* = 0.02) and the DNA polymerase cofactor PCNA (*p* = 0.04), belonging to the cellular component organization or biogenesis family, for instance (**Figure 3C**). Among the DEPs, the helicases are affected (13/77, 17%) but diversely, 8 being upregulated (DHX8, DDX10, ERCC3, DDX60 and DDX39B, *p* = 1.0e-6; DDX24, ATRX and DDX1, *p* < 0.03) and 5 being downregulated (DDX58 and ERCC6L, *p* = 1.0e-6; MCM6, EIF4A1 and DHX15, *p* < 0.04) (**Figure 3C**). Similarly, DNA repair proteins are diversely dysregulated (both double-strand break repair (7/52, 13%) and recombinational repair (5/30, 17%), but not in a statistically significant manner (*p* ≥ 0.5). These results indicated that the modulation of the proteome triggered by PDS is quite a comprehensive but also complex and multi-level process.

**Figure 3.**
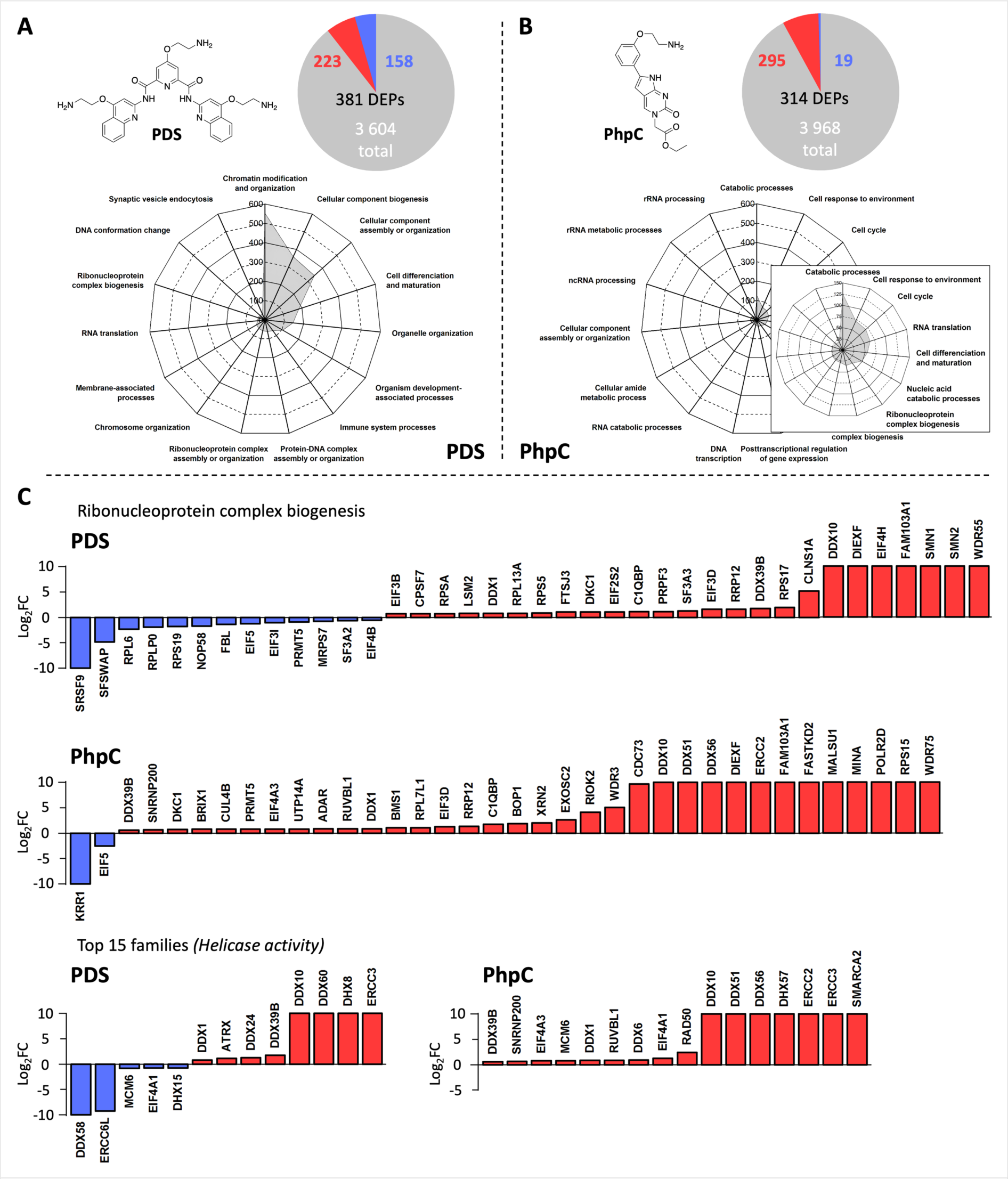
**A**. Differentially expressed proteins (DEPs) in upon incubation with PDS (pie chart) and dysregulated biological pathways (radar plot). **B**. DEPs upon incubation with PhpC (pie chart) and dysregulated biological pathways (radar plot). **C**. Examples of transcripts downregulated (blue) or upregulated (red) upon incubation with either PDS or PhpC belonging to families of dysregulated cellular pathways (see the supplementary data).

Compared to RNA-seq results, proteomic changes triggered by PhpC are much more significant: we identified 314 DEPs (7.9%) out of 3,968 proteins with measured expression. Among 314 DEPs, 295 proteins (94%) were shown to be upregulated, whereas only 19 proteins (6%) were downregulated (**Figure 3B** and Supplementary File 2). These results confirm that PhpC has more impact on astrocytes’ translational *versus* transcriptional activity. These modifications include 337 GO terms, 70 upstream regulators related to 16 pathways. Among them, the most notable pathways include base excision repair (4/16, 25%, including FEN1, *p* = 1.0e-6) for instance. The cellular processes affected are P-body assembly (4/10, 40%, including DDX6, *p* = 0.05), DNA transcription (8/38, 21%, including DDX39B, *p* = 0.03), regulation of RNA stability (13/77, 17%, including FUS, *p* = 1.0e-6, RNA stability), DNA duplex unwinding (6/40, 15%, including DDX1, *p* = 0.03, DNA repair), regulation of translation (26/188, 14%, including EIF5, *p* = 1.0e-6, RNA translation), among others **(Figure 3C)**. As above, among the DEPs, both helicases (with the upregulation of 16/83 helicases (19%), 8 being statistically relevant, DDX10, DDX51, DDX56, ERCC2 and ERCC3, DHX57, SMARCA2 and MCM6, *p* = 1.0e-6) (**Figure 3C**) and DNA damage binding proteins rank high (with the upregulation of 6/24 proteins (25%), 4 being statistically relevant, FEN1, ERCC2, ERCC3 and RAD23B, *p* = 1.0e-6).

Next, we investigated the common proteins between PDS and PhpC datasets. The expression of 107 proteins was found to be dysregulated by the 2 compounds (**Figure 4A** and Supplementary File 2), including 92 upregulated proteins (86%) in both instances (hereafter called PDS^+^/PhpC^+^), 5 downregulated (5%) in both cases (PDS^-^/PhpC^-^) and 10 differently expressed (9%), with 9 PDS^-^/PhpC^+^ and only 1 PDS^+^/PhpC^-^ (NUCKS1, *p* < 0.02). We found 33 biological processes dysregulated both by PDS and PhpC, including the significantly affected regulation of protein binding (22-27% of dysregulated proteins out of >30 proteins, *p* = 5.0e-3, protein binding family), ribonucleoprotein complex biogenesis (14-16% of >200 proteins, *p* = 3.8e-4, ribonucleoprotein complex biogenesis family) and cellular component organization (8-12% of >1,700 proteins, *p* = 7.0e-4, cellular component assembly or organization family) (**Figure 4B**), which are all related to the translation machinery.

**Figure 4.**
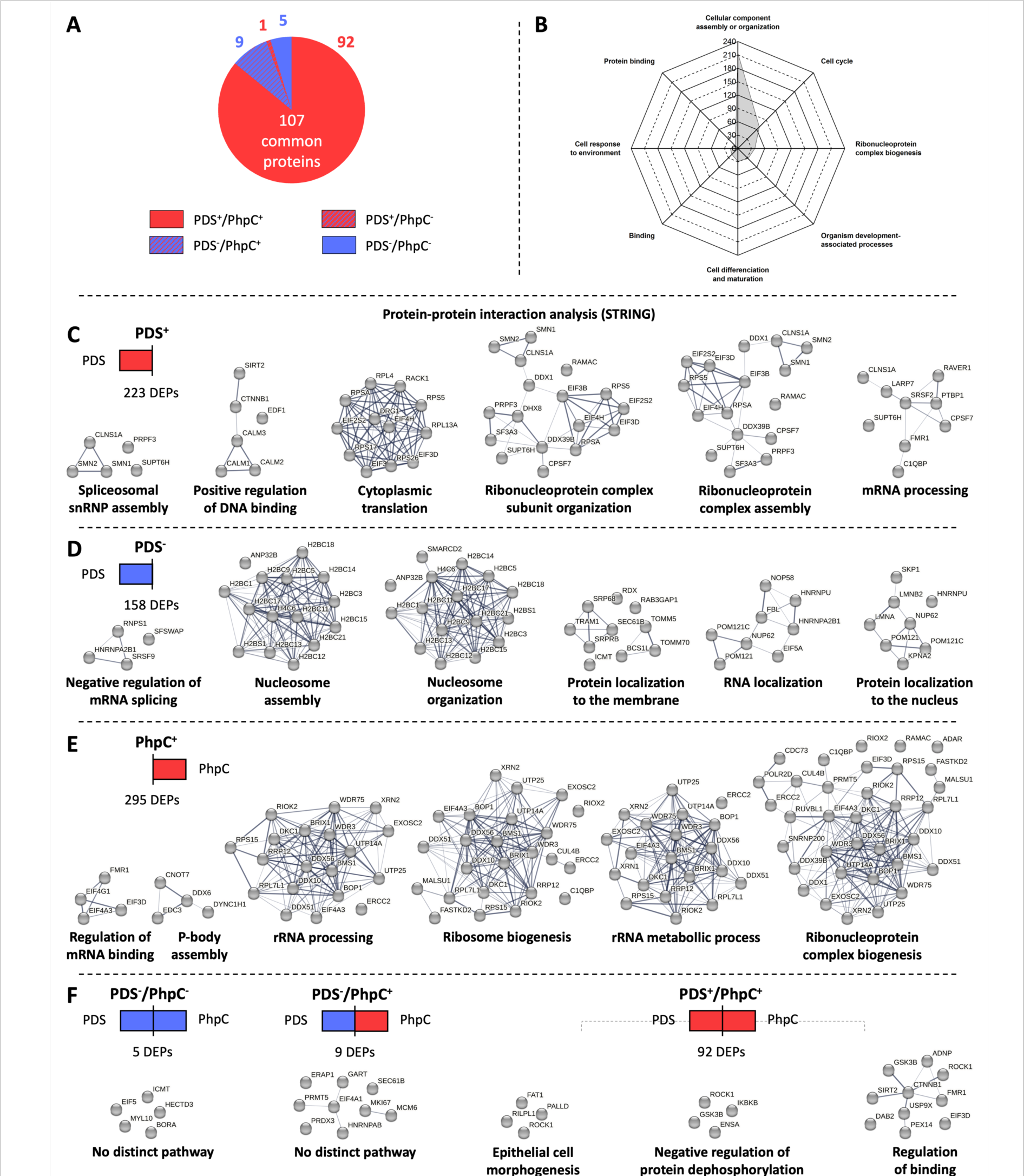
**A-B**. Common DEPs upon incubation with either PDS or PhpC (pie chart, **A**) and corresponding dysregulated biological pathways (radar plot, **B**). **C-F**. Examples of enrichment maps showing STRING connections (gene sets: node; gene presence: edge) for PDS^+^ (**C**), PDS^-^ (**D**), PhpC^+^ (**E**) and PDS^-^/PhpC^-^, PDS^-^/PhpC^+^ and PDS^+^/PhpC^+^ (**F**) proteins, gathered in families of dysregulated cellular pathways (see the supplementary data).

To gain more accurate insights into the cellular circuitries dysregulated by PDS, PhpC, or both, we performed a STRING analysis^63^ to predict functional association, quantified by a strength (*s*) score, considered to be high confidence when *s* ≥ 0.70. We found that the 223 PDS^+^ proteins are involved in RNA transactions (**Figure 4C** and Supplementary File 3), including cytoplasmic translation (12 proteins out of 123 (12/123), 10%, *s* = 0.94) and regulation of mRNA processing (9/140, 6%, *s* = 0.76), in ribonucleoprotein complex subunit assembly (16/203, 8%, *s* = 0.84) and organization (17/211, 8%, *s* = 0.85) and DNA binding (6/58, 3%, *s* = 0.96). In contrast, the 158 PDS^-^ proteins (**Figure 4D**) are primarily involved in cell organization (*e.g*., RNA localization (8/180, 4%, *s* = 0.79) and protein localization to the nucleus (8/195, 4%, *s* = 0.75) or membrane (10/219, 5%, *s* = 0.80)) and chromatin organization (*e.g*., nucleosome organization (16/162, 10%, *s* = 1.13) and assembly (15/124, 12%, *s* = 1.22)). Next, we focused on the 295 PhpC^+^ proteins (**Figure 4E**) and found that they are also involved in RNA transactions including rRNA processing (19/220, 9%, *s* = 0.76) and metabolic process (20/250, 8%, *s* = 0.73), ribosome (24/299, 8%, *s* = 0.73) and ribonucleoprotein complex biogenesis (34/449, 8%, *s* = 0.71). Notably, proteins involved in P-body assembly (4/14, 29%, *s* = 1.28) rank high, contributing to RNA transactions (degradation and silencing).^64^ In sharp contrast, the 19 PhpC^-^ proteins do not belong to any identifiable network, as do the 5 PDS^-^/PhpC^-^, 9 PDS^-^/PhpC^+^, and 1 PDS^+^/PhpC^-^ proteins (**Figure 4F**), likely due to their low number. We found that the 92 PDS^+^/PhpC^+^ proteins (**Figure 4F**) primarily intervene in protein dephosphorylation (4/34, 12%, *s* = 1.4), epithelial cell morphogenesis (4/34, 12%, *s* = 1.4) and more generally binding interactions (10/375, 3%, *s* = 0.76), but also in cellular component biogenesis (11/500, 2%), localization (29/2677, 1%) and organization (both cellular (48/5639, 0.9%) and organelle organization (37/3470, 1%)), although in a non-significant manner (*s* < 0.7). Interestingly, the molecular components that are most affected by both PDS and PhpC are associated to stress granule (7/84, 8%, *s* = 1.25) and polysome formation (5/65, 8%, *s* = 1.22), fully in line with the pivotal role play by G4s in response to cellular stress and in translation. ^65–67^ It is thus worth noting that both PDS and PhpC strongly impact RNA metabolism, although through different angles. PDS triggers a scatter-gun type of damage, affecting most if not all cellular circuities, which is in line with the RNA-seq analysis: indeed, the 354 PDS^-^ transcripts were found to belong to dozens of different cellular pathways (including DNA repair (62/497, 13%), replication (51/203, 25%) and unwinding (25/89, 28%), etc.) while, and quite surprisingly, the 162 PDS^+^ transcript do not belong to any identifiable network. This suggests that the cellular effects of PDS originate in its ability to downregulate cellular processes, acting at both transcriptional and translational levels (*i.e.*, on both DNA and RNA G4s). In sharp contrast, PhpC has a narrower spectrum of action, acting solely at the translational level (*i.e.*, on RNA G4s) and solely as an activating agent, positively regulating RNA processes, stress responses, and cellular internal organization.

Altogether, the results collected through both RNA-seq and proteomic analyses indicate that even if PDS and PhpC share the same targets (G4s), their cellular responses are strongly different, which is fully in line with the type of interaction they intend to have with G4s (stabilization *versus* destabilization). However, the G4 involvement in the cellular pathways and biological processes described above remained, at this stage, speculative. It was already established for some of them in the literature (*e.g*., the G4-forming sequence found in the promoter region of PCNA and BLM, in the 5’-untranslated region (UTR) of FEN1 mRNA (*vide infra*) for instance)^68, 69^ but the existence of G4s related to most of the aforementioned DEGs/DEPs described above remained to be demonstrated. To investigate this in detail, we evaluated the G4 content of the mRNA coding the DEPs using G4Hunter.^12^ From the 107 proteins found to be dysregulated by the 2 compounds, we selected only the DEPs whose statistical significance was established upon both PDS- and PhpC-treatment (*p* ≤ 0.01). This selection led to 58 candidates (**Figure 5A** and Supplementary File 4), including 4 PDS^-^/PhpC^-^ (*i.e.*, EIF5, HECTD3, ICMT, MYL10), 4 PDS^-^/PhpC^+^ (*i.e.*, EIF4A1, MKI67, PRMT5, SEC61B) and 50 PDS^+^/PhpC^+^ proteins (*e.g.*, DDX10, ERCC3, etc.). The corresponding mRNA sequences, obtained from the National Center for Biotechnology Information (NCBI) using the reference genome assembly GRCh38.p14 (GCF_000001405.40), were computed to identify their putative quadruplex sequences (PQS). We selected a rather low detection threshold (= 1.0 *versus* the classically used 1.5) for two reasons: first, we calibrated our investigations with previously established RNA G4-forming sequences and found that they display low scores such as FEN1 (see above, ^5’^GCAG_2_CUCAG_2_CUGCUG_4_C_2_GAGCAG_2_AG_2_UG_2_^3’^, +449/+511, score = 1.3));^68^ second, it has been reported that G4 ligands (notably PDS) preferentially stabilizes weaker G4s (that is, G4s with an average score centered on *ca*. 1.0) in human cells.^70^ Using these conditions, we found that the 58 studied mRNA contain at least 1 PQS, *i.e.*, 4 to 13 PQSs for PDS^-^/PhpC^-^ DEPs (score ≤ 2.00), 1 to 27 for PDS^-^/PhpC^+^ (score ≤ 2.35), and 1 to 58 for PDS^+^/PhpC^+^ (score ≤ 2.75), which represents a high G4-content, with an average of 9.5 PQS/transcript for PDS^-^/PhpC^-^, 10.5 for PDS^-^/PhpC^+^ and 10.9 for PDS^+^/PhpC^+^ (**Figure 5B** and Supplementary File 4).

**Figure 5.**
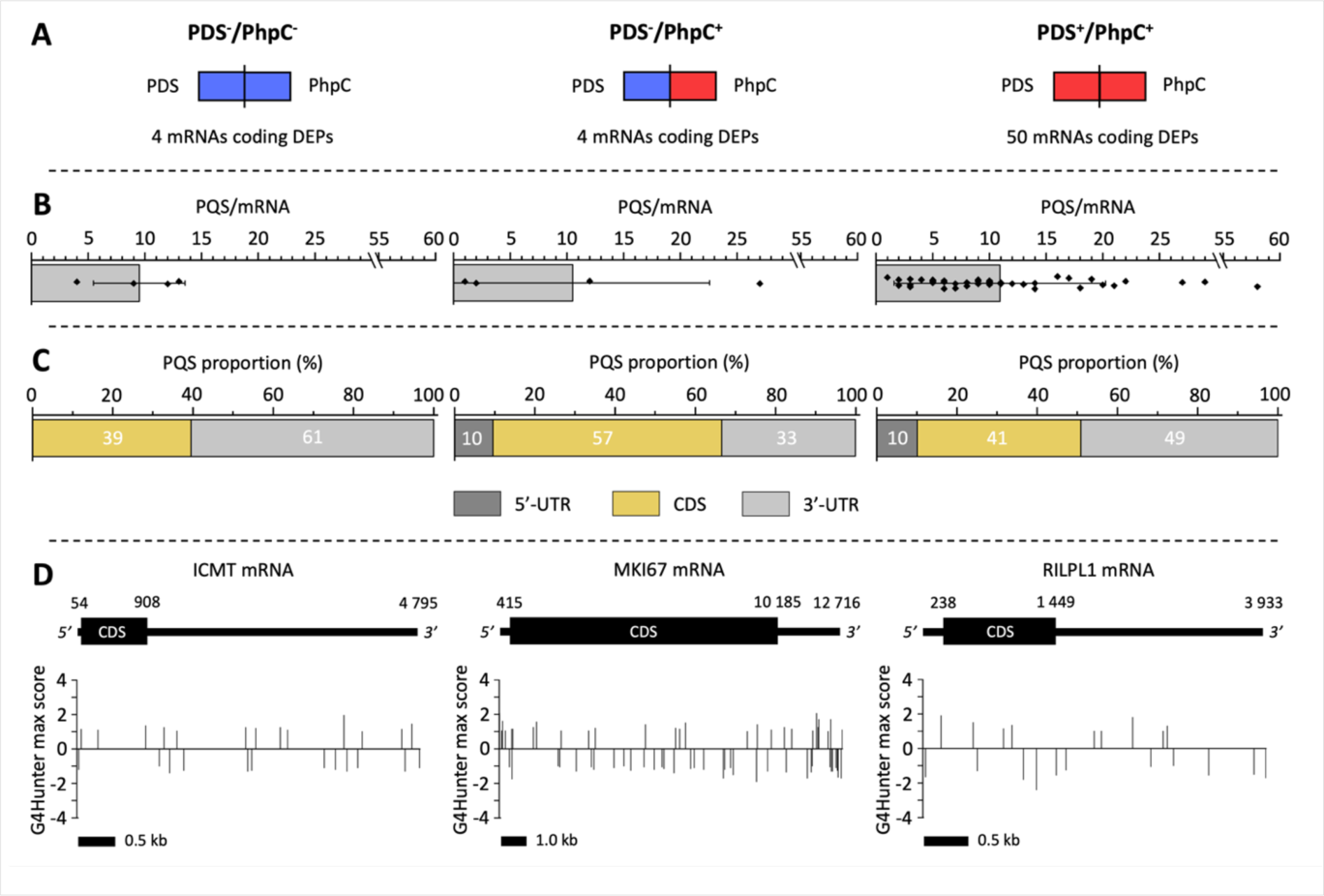
G4 content estimation within mRNA coding 58 significantly DEPs (**A**, 4 PDS^-^/PhpC^-^, 4 PDS^-^/PhpC^+^ and 50 PDS^+^/PhpC^+^) using G4hunter, with the distribution of PQS *per* mRNA (**B**) and across the various sections of mRNA (**C**), and the visualization of this distribution for a selection of mRNA belonging to each category (**D**).

To go a step further, the localization of these PQSs across the different mRNA sections (5’-UTR, coding sequence (CDS) and 3’-UTR) was investigated (**Figure 5C, D**): in the PDS^-^/PhpC^-^ transcripts, PQS are mainly located in the 3’-UTR (23 PQSs out of 38 (23/38), 61%) *versus* CDS (15/38, 39%); in the PDS^-^/PhpC^+^ transcripts, the PQSs are mainly located in CDS (24/42, 57%) *versus* 5’-UTR (4/42, 10%) and 3’-UTR (14/42, 33%); in the PDS^+^/PhpC^+^ transcripts, the PQSs are mainly located in the 3’-UTR (268/546, 49%) *versus* CDS (223/546, 41%) and 5’-UTR (55/546, 10%). This might have consequences in the regulation of mRNA expression: for instance, the transcripts whose expression is upregulated by PhpC tend to have a high G4-content in their CDS (57 *versus* 40% for PhpC^-^), which is in line with the known ability of these G4s to stall polymerases; they also have PQS in their 5’-UTR (10 *versus* 0% for PhpC^-^), which is also consistent with the ability of these G4s to hamper proper mRNA scanning;^71^ conversely, those downregulated by PhpC display a high PQS content in 3’-UTR (61 *versus* 33% for PhpC^+^), which is in agreement with the influence of these G4s on mRNA stability.^72^ Altogether, these *in silico* investigations demonstrated the G4-richness of identified DEGs/DEPs and confirmed their reliability as targets for both PDS and PhpC.

## Discussion

We performed a comprehensive study by systematically comparing the effects triggered upon either G4 stabilization by PDS or G4 destabilization by PhpC at both transcriptome- and proteome-wide scales in primary human astrocytes. Our results confirm that, just like in many other cell types,^55, 73–76^ PDS is a quite active, pan-G4 ligand that affects a broad variety of cellular processes and that its toxicity primarily originates in its inhibitory properties, with the downregulation of 354 transcripts (RNA-seq) and 158 proteins (mass spectrometry), which affects many different pathways mostly related to RNA transactions. In contrast, PhpC modulates the G4 landscapes in a more focused manner, altering only the translational activity in astrocytes (likely because of a preferential interaction with RNA G4s, previously demonstrated by G4RP) with the upregulation of 295 proteins mostly involved in RNA transactions, thus mirroring the effects of PDS.

From the G4 point of view, the 107 proteins dysregulated by both PDS and PhpC offer a unique and dazzling display of the role that G4s play in human astrocytes. It is important to stress that G4 folding and unfolding do not necessarily correspond to inhibition and activation of cellular processes, respectively. Indeed, depending on the localization of the G4-forming sequence within the gene/transcript (UTR or CDS), G4s can promote (behaving as docking sites, for instance)^77–79^ or hamper (behaving as an impediment to polymerase motion, for instance)^80–82^ the related transactions. Therefore, both PDS^-^ and PDS^+^, and accordingly PhpC^-^ and PhpC^+^ can be interpreted to cast a light on the involvement of G4s in cellular biology. Among this set of dysregulated proteins, both iPathwayGuide and STRING analyses point toward key G4 roles in the regulation of stress response and protein production and localization, thus confirming the functional prevalence of G4-associated pathways in astrocytes and the relevance of the strategies aiming at targeting G4s to harness the cells’ operational pattern.

From a molecular point of view, our results confirm that PDS causes casualties in a rather indiscriminate manner, up- and downregulating gene expression at both transcriptional (516 DEGs, 31% up, 69% down) and translational steps (381 DEPs, 58% up, 42% down). PhpC behaves in a radically different manner, upregulating gene expression at the translational step only (9 DEGs only *versus* 314 DEPs, 94% up, 6% down). PDS and PhpC might not interact with the same G4s, even within the same gene/transcript, nor with the same affinity; in spite of these differences, we found that PhpC triggers effects opposite to PDS, favoring sheltering effects while PDS favors inhibitory effects. PhpC could, therefore, be considered as a senolytic drug that may find many applications in designing strategies to prevent G4-associated molecular and genetic dysfunctions associated with age-related neurological disorders.

## Materials and methods

### Isolation and culture of primary human astrocytes

In this study, we collected epileptic brain tissue resection from the frontal lobe of a 32-year-old male patient, maintaining sterility throughout the process. The patient had no known pathological conditions or co-morbidities, and ethical standards, including obtaining informed consent, were observed throughout the handling of human tissue sample. The brain sample was subjected to a 30-second sterilization step using 20% Betadine solution, followed by thorough rinsing with 1X PBS. Subsequently, the tissue was finely diced into 1mm-sized pieces on a sterile petri dish, and these explants were then transferred to PDL-coated plates and air-dried for 10 minutes. The explants were cultures in DMEM growth medium supplemented with 10% FBS, as well as antibiotics, penicillin and streptomycin. After culturing them for 2-3 weeks, astrocytes were isolated from the explants and continued their culture in DMEM growth medium enriched with recombinant human TGF-α and human NRG1-β1 growth factors, promoting their maturation and differentiation until 3-4 passages.

### *In Vitro* maturation of human astrocytes and small molecule treatment for RNA-sequencing and mass spectrometry analysis

Human astrocytes, isolated from the explants, were cultured *in vitro* in DMEM media supplemented with FBS and growth factors for a minimum of 3-4 passages, ensuring their maturation and differentiation. Astrocytes were cultured in PDL-coated T25 flasks at a seeding density of approximately 2 million cells per flask. Cells were treated with PDS (2 µM) or PhpC (20 µM) for 24 hrs. The following day, for RNA-sequencing, the cells were lysed in 1 mL of RLT buffer obtained from Qiagen, and for Mass spectrometry, cells were collected in 1X PBS. Cells were rapidly frozen and stored at -80°C until further analysis.

### Immunocytochemistry

Cultured human astrocytes were grown on Geltrex-coated chambered slides and subjected to immunofluorescence staining. After a preliminary wash with 1X Phosphate-Buffered Saline (PBS), the cells were fixed using 4% paraformaldehyde (PFA) for 10 minutes at room temperature, followed by PBS washes. Subsequent permeabilization and blocking were done using 5% normal goat serum, 1% Bovine Serum Albumin (BSA), and 0.3% Triton-X-100 for 45 minutes at room temperature. The astrocytes were then incubated with a primary antibody prepared in the blocking solution overnight at 4°C, washed thrice with PBS for 5 minutes each, and subsequently incubated with a fluorophore-tagged secondary antibody for 1 hour at room temperature. The slides were mounted with a DAPI-containing mounting medium, enabling visualization using fluorescence microscopy.

### Fluorescence microscopy and image analyses

Fixed cells were imaged using a Nikon A1R Confocal Laser Microscope, employing 20X, 40X, and 63X objectives. To ensure uniformity and comparability across all samples, imaging parameters, including light intensity, gain, and other relevant settings, were maintained constant throughout the imaging process. Subsequently, the acquired images were analyzed and quantified using ImageJ/Fiji software, enabling standardized measurements and evaluations of the experimental data.

### Sample preparation, library preparation, mRNA sequencing, and data analysis workflow

RNA isolation was performed using the RNeasy Mini kit (Qiagen) following the manufacturer’s guidelines, with elution on a 30 µL volume. Subsequent library preparation was carried out adhering to the standard QIAseq stranded mRNA Kit (Qiagen) protocol. Starting with 500 ng of initial material, mRNA was enriched and heat fragmented. Following the first and second strand synthesis, complementary DNA underwent end-repair and 3’ adenylation. Sequencing adapters were then ligated to the overhangs, and molecules carrying adapters were enriched through 13 cycles of PCR and purified using bead-based cleanup. Library quality control was performed using capillary electrophoresis (Tape D1000), and high-quality libraries were pooled based on equimolar concentrations. The concentration of the library pool was quantified using qPCR, and this optimal concentration was utilized to generate clusters on the surface of a flow cell. Sequencing was conducted on a Nextseq instrument (Illumina Inc.) according to the manufacturer’s instructions, producing paired-end reads (2×75, 2×10). Primary data analysis was carried out using CLC Genomics Server 22.0.02. Initial data processing involved trimming to remove potential read-through adapter sequences at the 3’end, followed by quality score-based trimming and handling of reads containing ambiguous nucleotides (up to 2 per read). The trimmed reads were first mapped to the Human ribosomal RNA (rRNA) repetitive unit to assess the rRNA content in the samples. Subsequently, all reads were mapped to the Human genome GRCh38 with ENSEMBL GRCh38 version 98 annotation. For differential expression analysis, the ‘Empirical analysis of DGE’ algorithm within the CLC Genomics Workbench 22.0.2 was employed with default settings, utilizing the ‘Exact Test’ from the EdgeR Bioconductor package for two-group comparisons. Only genes with at least 10 counts summed across all samples were considered for unsupervised analysis. A variance stabilizing transformation was applied to the raw count matrix using the R package DESeq2 version 1.28.1. 500 genes with the highest variance were used for the principal component analysis, and the variance values were calculated agnostically to pre-defined groups (blind=TRUE). Finally, hierarchical clustering was performed using the top 35 genes exhibiting the highest variance across samples.

### Sample preparation, library preparation, and mass spectrometry

Astrocytic cultures were lysed with 50 mM ammonium bicarbonate and 1 mM calcium chloride with sonication (3 mins, 30 Amp, 30-second pulse, a 1-min rest at 4°C). The protein concentration of the lysate was measured using the Bradford method, and 25 µg of lysate were digested with 0.5 µg of trypsin for 12 hours at 37°C. The concentration of tryptic peptides was measured using the Pierce Quantitative Peptide Assays kit (ThermoFisher Scientific cat# 23275), and 25 µg of peptides were separated on a home-made reverse-phase C18 column in a pipet tip. Peptides were eluted and separated into fifteen fractions using a stepwise gradient of increasing acetonitrile (2, 4, 6, 8, 10, 12, 14, 16, 18, 20, 22, 24, 26, 28 and 35% acetonitrile) at pH 10. Subsequently, these fractions were combined into five groups (2+12+22, 4+14+24, 6+16+26, 8+18+28, 10+20+35) and vacuum-dried. The dried peptide samples were resuspended in a solution of 5% methanol and 0.1% formic acid in water, then analyzed using Orbitrap Fusion mass spectrometers (Thermo Fisher Scientific) coupled with an Easy-nLC 1000 nanoflow LC system (Thermo Fisher Scientific). The analysis utilized an in-house trap column packed with 1.9 μm Reprosil-Pur Basic C18 beads (2 cm × 100 μm) and a 5 cm × 150 μm capillary separation column packed with 1.9 μm Reprosil-Pur Basic C18 beads. The separation was carried out using a 75-minute discontinuous gradient of 4–26% acetonitrile and 0.1% formic acid at a flow rate of 800 nl/min. The instrument was operated under the control of Xcalibur software version 4.1 (Thermo Fisher Scientific) in data-dependent mode, acquiring fragmentation spectra of the top 3 seconds. Parent mass spectrum was acquired in the Orbitrap with a full MS range of 300–1400 m/z at the resolution of 120,000. Higher-energy collisional dissociation (HCD) fragmented MS/MS spectrum was acquired in ion-trap with rapid scan mode. The MS/MS spectra were searched against the target-decoy mouse RefSeq database (release March 23, 2020, containing 62,232 entries) in Proteome Discoverer 2.1 interface (Thermo Fisher) with Mascot algorithm (Mascot 2.4, Matrix Science). The precursor mass tolerance of 20 ppm and fragment mass tolerance of 0.5 Da was allowed. Two maximum missed cleavage and dynamic modifications of acetylation of N-term and oxidation of methionine were allowed. Assigned peptides were filtered with a 1% false discovery rate (FDR) using Percolator validation based on q-value. The Peptide Spectrum Matches (PSMs) output from PD2.1 was used to group peptides onto gene level using the ‘gpGrouper’ algorithm56. An in-housed program, gpGrouper, uses a universal peptide grouping logic to accurately allocate and provide MS1-based quantification across multiple gene products. Gene-protein products (GPs) quantification was performed using the label-free, intensity-based absolute quantification (iBAQ) approach and then normalized to FOT (a fraction of the total protein iBAQ amount per experiment). FOT was defined as an individual protein’s iBAQ divided by the total iBAQ of all identified proteins within one experiment.

### RNA-sequencing and Mass spectrometry data analysis

The RNA sequencing and mass spectrometry data were subjected to analysis using iPathwayGuide, a tool provided by Adviata Bioinformatics. The analysis was performed within the context of pathways sourced from the KEGG (Kyoto Encyclopedia of Genes and Genomes) database, as well as gene ontologies obtained from the GO Consortium database. The analysis integrated these data sources to construct underlying topologies, which represent the network of genes and proteins and their directional interactions. The topologies were derived from the KEGG database utilizing iPathwayGuide, providing valuable insights into the biological pathways and processes influenced by the differential gene expression patterns identified through RNA sequencing and mass spectrometry.

### String analysis

Functional network analysis was performed using STRING (https://string-db.org/) for 6 families of differently expressed proteins (DEPs) extracted from the iPathwayGuide (Advaita Bioinformatics) protein lists provided in the Supplementary file 3 (*i.e*., 223 PDS^+^, 295 PhpC^+^, 5 PDS^-^/PhpC^-^, 9 PDS^-^/PhpC^+^ and 92 PDS^+^/PhpC^+^ proteins) using *Homo sapiens* organism, full STRING network, evidence network edges, medium confidence interaction score (0.400) and confidence strength (*s*) ≥ 0.70 within Biological Process (gene Ontology) as outputs.

### G4 motif analysis using G4Hunter

G4Hunter scores were computed using specific parameters: information (length in base, genomic subdomains (UTRs and CDS) positions) and FASTA files of mRNAs (real variant 1, regardless of the number of real and predicted mRNA or ncRNA variants) coding DEPs were first extracted from the reference genome assembly GRCh38.p14 (GCF_000001405.40) *via* the NCBI database and then processed through the dedicated G4Hunter website (https://g4hunter.shinyapps.io/G4HunterMultiFastaSeeker/) with a window of 20 and a minimal score (threshold) of 1.0. Each ‘hit’ (or putative G-quadruplex sequence, PQS) is characterized by its genomic coordinates and a ‘max_score’ reflecting the score of the highest G4Hunter within this window. These data are available in the Supplementary file 4. Number of PQS per genomic subdomains for each significant common DEPs groups (PDS^-^/PhpC^-^, PDS^-^/PhpC^+^ and PDS^+^/PhpC^+^) was gathered manually, analyzed with Excel (Microsoft Corp.) and OriginPro (OriginLab Corporation).

### Ethical approval

Human brain tissue samples were obtained and handled in strict adherence to the guidelines and regulations established by the University of Texas, McGovern Medical School at Houston. Ethical considerations and prior consent from the patients were obtained and approved to enable research involving the use of their brain tissue. All experimental protocols underwent approval by the University of Texas, McGovern Medical School at Houston, and the research procedures were conducted in full compliance with the approved guidelines, ensuring ethical and scientific standards throughout the study.

### Conflict of interest

The authors declare no conflict of interest with the content of the article.

## Acknowledgments

We thank members of the A. S. T. laboratory and the BRAINS laboratory for useful discussions. We also thank Suresh Kumar (bioinformatics scientist, Advaita Bioinformatics) for helpful discussions. Danielle Guillory provided administrative assistance. This work was supported by the Glenn Foundation and the American Federation for Aging Research (grant ID: BIG21042) to A. S. T., and by the *Ligue nationale contre le cancer* (grant ID: AAPEAC2022.LCC/VG) and the *Centre national de la recherche scientifique* (CNRS) to D.M. We also thank Laurent Lacroix for helpful discussions about G4Hunter.

## Notes

### Competing Interest Statement

The authors have declared no competing interest.

## References

1. Pauling, L. & Beadle, G.W. Chemical Biology. Eng. Sci. 17, 9–13 (1954).

2. Schreiber, S.L. Small molecules: the missing link in the central dogma. Nat. Chem. Biol. 1, 64 (2005).

3. Altmann, K.-H. et al. The state of the art of chemical biology. ChemBioChem 10, 16–29 (2009).

4. Spiegel, J., Adhikari, S. & Balasubramanian, S. The structure and function of DNA G-quadruplexes. Trends Chem. 2, 123–136 (2020).

5. Dumas, L., Herviou, P., Dassi, E., Cammas, A. & Millevoi, S. G-Quadruplexes in RNA Biology: Recent Advances and Future Directions. Trends Biochem. Sci. 46, 270–283 (2021).

6. Varshney, D., Spiegel, J., Zyner, K., Tannahill, D. & Balasubramanian, S. The regulation and functions of DNA and RNA G-quadruplexes. Nat. Rev. Mol. Cell Biol. 21, 459–474 (2020).

7. Lyu, K., Chow, E.Y.-C., Mou, X., Chan, T.-F. & Kwok, Chun K. RNA G-quadruplexes (rG4s): genomics and biological functions. Nucleic Acids Res. 49, 5426–5450 (2021).

8. Raguseo, F., Chowdhury, S., Minard, A. & Di Antonio, M. Chemical-biology approaches to probe DNA and RNA G-quadruplex structures in the genome. Chem. Commun. 56, 1317–1324 (2020).

9. Stefan, L. & Monchaud, D. Applications of guanine quartets in nanotechnology and chemical biology. Nat. Rev. Chem. 3, 650–668 (2019).

10. Cammas, A. & Millevoi, S. RNA G-quadruplexes: emerging mechanisms in disease. Nucleic Acids Res. 45, 1584–1595 (2017).

11. Ganser, L.R., Kelly, M.L., Herschlag, D. & Al-Hashimi, H.M. The roles of structural dynamics in the cellular functions of RNAs. Nat. Rev. Mol. Cell Biol., 474–489 (2019).

12. Bedrat, A., Lacroix, L. & Mergny, J.-L. Re-evaluation of G-quadruplex propensity with G4Hunter. Nucleic Acids Res. 44, 1746–1759 (2016).

13. Vannutelli, A., Belhamiti, S., Garant, J.-M., Ouangraoua, A. & Perreault, J.-P. Where are G-quadruplexes located in the human transcriptome? NAR Genomics and Bioinformatics 2 (2020).

14. Meier-Stephenson, V. G4-quadruplex-binding proteins: review and insights into selectivity. Biophys. Rev. 14, 635–654 (2022).

15. Brosh, R.M. & Matson, S.W. History of DNA Helicases. Genes 11, 255 (2020).

16. Lerner, L.K. & Sale, J.E. Replication of G quadruplex DNA. Genes 10, 95 (2019).

17. Lejault, P., Mitteaux, J., Rota Sperti, F. & Monchaud, D. How to untie G-quadruplex knots and why? Cell Chem. Biol. 28, 436–455 (2021).

18. Antcliff, A., McCullough, L.D. & Tsvetkov, A.S. G-Quadruplexes and the DNA/RNA helicase DHX36 in health, disease, and aging. Aging (Albany NY) 13, 25578 (2021).

19. Vijay Kumar, M.J., Morales, R. & Tsvetkov, A.S. G-quadruplexes and associated proteins in aging and Alzheimer’s disease. Front Aging 4, 1164057 (2023).

20. Neidle, S. Quadruplex Nucleic Acids as Novel Therapeutic Targets. J. Med. Chem. 59, 5987–6011 (2016).

21. Monchaud, D. in Annual Reports in Medicinal Chemistry, Vol. 54. (ed. S. Neidle) 133–160 (Academic Press, 2020).

22. Robinson, J., Raguseo, F., Nuccio, S.P., Liano, D. & Di Antonio, M. DNA G-quadruplex structures: more than simple roadblocks to transcription? Nucleic Acids Res. 49, 8419–8431 (2021).

23. Kosiol, N., Juranek, S., Brossart, P., Heine, A. & Paeschke, K. G-quadruplexes: A promising target for cancer therapy. Mol. Cancer 20, 40 (2021).

24. del Mundo, I.M., Vasquez, K.M. & Wang, G. Modulation of DNA structure formation using small molecules. Biochim. Biophys. Acta - Mol. Cell Res., 118539 (2019).

25. Weisman-Shomer, P. et al. The cationic porphyrin TMPyP4 destabilizes the tetraplex form of the fragile X syndrome expanded sequence d (CGG) n. Nucleic Acids Res. 31, 3963–3970 (2003).

26. Waller, Z.A.E., Sewitz, S.A., Hsu, S.-T.D. & Balasubramanian, S. A Small Molecule That Disrupts G-Quadruplex DNA Structure and Enhances Gene Expression. J. Am. Chem. Soc. 131, 12628–12633 (2009).

27. Zamiri, B., Reddy, K., Macgregor, R.B. & Pearson, C.E. TMPyP4 porphyrin distorts RNA G-quadruplex structures of the disease-associated r (GGGGCC) n repeat of the C9orf72 gene and blocks interaction of RNA-binding proteins. J. Biol. Chem. 289, 4653–4659 (2014).

28. Kaluzhny, D. et al. Disordering of Human Telomeric G-Quadruplex with Novel Antiproliferative Anthrathiophenedione. PLoS One 6, e27151 (2011).

29. Mitteaux, J. et al. Identifying G-Quadruplex-DNA-Disrupting Small Molecules. J. Am. Chem. Soc. 143, 12567–12577 (2021).

30. Chowdhury, S., Wang, J., Nuccio, Sabrina P., Mao, H. & Di Antonio, M. Short LNA-modified oligonucleotide probes as efficient disruptors of DNA G-quadruplexes. Nucleic Acids Res. 50, 7247–7259 (2022).

31. Mitteaux, J. et al. PhpC modulates G-quadruplex-RNA landscapes in human cells. Chem. Commun. 60, 424–427 (2024).

32. Yang, S.Y., Monchaud, D. & Wong, J.M.Y. Global mapping of RNA G-quadruplexes (G4-RNAs) using G4RP-seq. Nat. Protoc. 17, 870–889 (2022).

33. Moruno-Manchon, J.F. et al. Small-molecule G-quadruplex stabilizers reveal a novel pathway of autophagy regulation in neurons. eLife 9, e52283 (2020).

34. Rodriguez, R. et al. A Novel Small Molecule That Alters Shelterin Integrity and Triggers a DNA-Damage Response at Telomeres. J. Am. Chem. Soc. 130, 15758–15758 (2008).

35. Moruno-Manchon, J.F. et al. The G-quadruplex DNA stabilizing drug pyridostatin promotes DNA damage and downregulates transcription of Brca1 in neurons. Aging (Albany NY) 9, 1957 (2017).

36. Tabor, N. et al. Differential responses of neurons, astrocytes, and microglia to G-quadruplex stabilization. Aging 13, 15917–15941 (2021).

37. Escarcega, R.D. et al. Pirh2-dependent DNA damage in neurons induced by the G-quadruplex ligand pyridostatin. J. Biol. Chem. 299, 105157 (2023).

38. Scheibye-Knudsen, M. et al. Cockayne syndrome group A and B proteins converge on transcription-linked resolution of non-B DNA. Proc. Natl. Acad. Sci. U. S. A. 113, 12502–12507 (2016).

39. Lee, J.-H., Kim, E.W., Croteau, D.L. & Bohr, V.A. Heterochromatin: an epigenetic point of view in aging. Experimental & Molecular Medicine 52, 1466–1474 (2020).

40. Sen, P., Shah, P.P., Nativio, R. & Berger, S.L. Epigenetic mechanisms of longevity and aging. Cell 166, 822–839 (2016).

41. Saul, D. & Kosinsky, R.L. Epigenetics of aging and aging-associated diseases. Int. J. Mol. Sci. 22, 401 (2021).

42. Tsurumi, A. & Li, W.X. Global heterochromatin loss: a unifying theory of aging? Epigenetics 7, 1–9 (2012).

43. Wang, E., Thombre, R., Shah, Y., Latanich, R. & Wang, J. G-Quadruplexes as pathogenic drivers in neurodegenerative disorders. Nucleic Acids Res. 49, 4816–4830 (2021).

44. Cave, J.W. & Willis, D.E. G-quadruplex regulation of neural gene expression. FEBS J. 289, 3284–3303 (2022).

45. Vijay Kumar, M., Morales, R. & Tsvetkov, A.S. G-quadruplexes and associated proteins in aging and Alzheimer’s disease. Frontiers in Aging 4, 1164057 (2023).

46. Jakel, S. & Dimou, L. Glial Cells and Their Function in the Adult Brain: A Journey through the History of Their Ablation. Front Cell Neurosci 11, 24 (2017).

47. Acioglu, C., Li, L. & Elkabes, S. Contribution of astrocytes to neuropathology of neurodegenerative diseases. Brain Res. 1758, 147291 (2021).

48. Brandebura, A.N., Paumier, A., Onur, T.S. & Allen, N.J. Astrocyte contribution to dysfunction, risk and progression in neurodegenerative disorders. Nat. Rev. Neurosci. 24, 23–39 (2023).

49. Phatnani, H. & Maniatis, T. Astrocytes in neurodegenerative disease. *Cold Spring Harbor Persp*. Biol. 7, a020628 (2015).

50. Palmer, A.L. & Ousman, S.S. Astrocytes and Aging. Frontiers in Aging Neuroscience 10 (2018).

51. Papadopoulos, M.C. & Verkman, A.S. Aquaporin water channels in the nervous system. Nat. Rev. Neurosci. 14, 265–277 (2013).

52. Kim, K. et al. Role of Excitatory Amino Acid Transporter-2 (EAAT2) and glutamate in neurodegeneration: Opportunities for developing novel therapeutics. J. Cell. Physiol. 226, 2484–2493 (2011).

53. Hol, E.M. & Pekny, M. Glial fibrillary acidic protein (GFAP) and the astrocyte intermediate filament system in diseases of the central nervous system. Curr. Opin. Cell Biol. 32, 121–130 (2015).

54. Chen, K.-Z. et al. Vimentin as a potential target for diverse nervous system diseases. Neural Regeneration Research 18, 969–975 (2023).

55. Miglietta, G., Russo, M., Duardo, R.C. & Capranico, G. G-quadruplex binders as cytostatic modulators of innate immune genes in cancer cells. Nucleic Acids Res. 49, 6673–6686 (2021).

56. Kanehisa, M., Furumichi, M., Tanabe, M., Sato, Y. & Morishima, K. KEGG: new perspectives on genomes, pathways, diseases and drugs. Nucleic Acids Res. 45, D353–D361 (2016).

57. Ashburner, M. et al. Gene Ontology: tool for the unification of biology. Nat. Genet. 25, 25–29 (2000).

58. Consortium, T.G.O. et al. The Gene Ontology knowledgebase in 2023. Genetics 224 (2023).

59. Ahsan, S. & Drăghici, S. Identifying Significantly Impacted Pathways and Putative Mechanisms with iPathwayGuide. Current Protocols in Bioinformatics 57, 7.15.11–17.15.30 (2017).

60. Rodriguez, R. et al. Small-molecule-induced DNA damage identifies alternative DNA structures in human genes. Nat. Chem. Biol. 8, 301–310 (2012).

61. Rota Sperti, F., et al. Click-Chemistry-Based Biomimetic Ligands Efficiently Capture G-Quadruplexes In Vitro and Help Localize Them at DNA Damage Sites in Human Cells. JACS Au 2, 1588–1595 (2022).

62. Blackford, A.N. & Jackson, S.P. ATM, ATR, and DNA-PK: The Trinity at the Heart of the DNA Damage Response. Mol. Cell 66, 801–817 (2017).

63. Szklarczyk, D. et al. STRING v10: protein–protein interaction networks, integrated over the tree of life. Nucleic Acids Res. 43, D447–D452 (2014).

64. Jia, L. et al. Decoding mRNA translatability and stability from the 5ʹ UTR. Nat. Struct. Mol. Biol. 27, 814–821 (2020).

65. Turner, M. et al. rG4detector, a novel RNA G-quadruplex predictor, uncovers their impact on stress granule formation. Nucleic Acids Res. 50, 11426–11441 (2022).

66. Kharel, P. & Ivanov, P. RNA G-quadruplexes and stress: emerging mechanisms and functions. Trends Cell Biol.

67. Rhodes, D. & Lipps, H.J. G-quadruplexes and their regulatory roles in biology. Nucleic Acids Res. 43, 8627–8637 (2015).

68. Ma, Y. et al. RNA G-Quadruplex within the 5’-UTR of FEN1 Regulates mRNA Stability under Oxidative Stress. Antioxidants 12, 276 (2023).

69. Fleming, A.M. et al. Human DNA Repair Genes Possess Potential G-Quadruplex Sequences in Their Promoters and 5ʹ-Untranslated Regions. Biochem. 57, 991–1002 (2018).

70. Esnault, C. et al. G4access identifies G-quadruplexes and their associations with open chromatin and imprinting control regions. Nat. Genet. 55, 1359–1369 (2023).

71. Leppek, K., Das, R. & Barna, M. Functional 5ʹ UTR mRNA structures in eukaryotic translation regulation and how to find them. Nat. Rev. Mol. Cell Biol. 19, 158–174 (2018).

72. Beaudoin, J.-D. & Perreault, J.-P. Exploring mRNA 3ʹ-UTR G-quadruplexes: evidence of roles in both alternative polyadenylation and mRNA shortening. Nucleic Acids Res. 41, 5898–5911 (2013).

73. Zimmer, J. et al. Targeting BRCA1 and BRCA2 Deficiencies with G-Quadruplex-Interacting Compounds. Mol. Cell 61, 449–460 (2016).

74. Zyner, K.G. et al. Genetic interactions of G-quadruplexes in humans. eLife 8, e46793 (2019).

75. Olivieri, M. et al. A Genetic Map of the Response to DNA Damage in Human Cells. Cell 182, 481–496.e421 (2020).

76. Bossaert, M. et al. Transcription-associated topoisomerase 2α (TOP2A) activity is a major effector of cytotoxicity induced by G-quadruplex ligands. eLife 10, e65184 (2021).

77. Spiegel, J. et al. G-quadruplexes are transcription factor binding hubs in human chromatin. Genome Biol. 22, 117 (2021).

78. Esain-Garcia, I. et al. G-quadruplex DNA structure is a positive regulator of MYC transcription. Proc. Natl. Acad. Sci. U. S. A. 121, e2320240121 (2024).

79. Song, J., Perreault, J.-P., Topisirovic, I. & Richard, S. RNA G-quadruplexes and their potential regulatory roles in translation. Translation 4, e1244031 (2016).

80. Murat, P., Guilbaud, G. & Sale, J.E. DNA polymerase stalling at structured DNA constrains the expansion of short tandem repeats. Genome Biol. 21, 209 (2020).

81. Lemmens, B., Van Schendel, R. & Tijsterman, M. Mutagenic consequences of a single G-quadruplex demonstrate mitotic inheritance of DNA replication fork barriers. Nat. Commun. 6, 8909 (2015).

82. Williams, S.L. et al. Replication-induced DNA secondary structures drive fork uncoupling and breakage. EMBO J. 42, e114334 (2023).

